# Microstructural correlates of white and gray matter in the healthy human brain: comparative analysis of diffusion biophysical models, inhomogeneous magnetization transfer, and macromolecular proton fraction

**DOI:** 10.64898/2025.12.06.692761

**Authors:** Andreea Hertanu, Tommaso Pavan, Quentin Uhl, Santiago Mezzano, Thorsten Feiweier, Ileana O. Jelescu

**Affiliations:** Dept. of Radiology, Lausanne University Hospital (CHUV) and University of Lausanne, Switzerland; Siemens Healthineers AG, Erlangen, Germany

**Keywords:** diffusion MRI, microstructure, brain, white/gray matter, Standard Model Imaging, Standard Model with Exchange, inhomogeneous Magnetization Transfer, Macromolecular Proton Fraction

## Abstract

Diffusion MRI (dMRI) and magnetization transfer (MT) rely on distinct biophysical principles and provide complementary insights into tissue microstructure. In this study, we investigated associations between microstructural metrics derived from the Standard Model (SMI) in white matter (WM) and the Standard Model with EXchange (SMEX/NEXI) in gray matter (GM) with two MT measures differing in specificity and sensitivity to myelination: the macromolecular proton fraction (MPF) and the inhomogeneous magnetization transfer ratio (ihMTR). Measurements were performed in WM and GM in ten healthy subjects scanned at 3T. In WM, the strongest significant association was observed between ihMTR and the axonal water fraction, consistent with higher myelination in regions of elevated axonal density and limited extra-axonal space. This correlation exceeded that of MPF, supporting the greater specificity of ihMTR to myelin. Interestingly, ihMTR displayed a gradient along the longitudinal axis of the corpus callosum, in agreement with previous histology measurements. Correlation between ihMTR and the extra-axonal perpendicular diffusivity (D_e_^⊥^), a putative myelination biomarker, was not significant, whereas MPF and D_e_^⊥^ exhibited a moderate significant positive correlation. Since a negative correlation is expected if reflecting myelination, these results suggest that in healthy tissue D_e_^⊥^ is mainly influenced by microstructural factors like fiber coherence and packing, the latter most likely affecting MPF but not ihMTR. In GM, ihMTR correlated significantly only with the exchange time (tₑₓ), confirming t_ex_ as a proxy for cell membrane permeability modulated by myelin. MPF correlated exclusively with the cell-process fraction (f), suggesting the latter is modulated by total (neuronal and glial) cell membrane density. Overall, findings underscore the complementary and concurring microstructural information captured by these metrics, highlighting their potential to disentangle distinct tissue mechanisms in both healthy and pathological conditions. Future studies incorporating ground-truth histology should validate the precise sensitivity of each metric to various microstructural tissue features.

## 1. INTRODUCTION

Biophysical models of diffusion MRI (dMRI) probe the fine-scale architecture of the central nervous system with greater specificity than conventional signal representation approaches such as diffusion tensor and kurtosis imaging (DTI/DKI)^1,2^. Within this framework, a family of multi-compartmental models collectively referred to as the Standard Model (SM)^3^ of diffusion in white matter (WM) has been developed to disentangle intra- and extra-axonal contributions to the dMRI signal. SM-derived parameters such as the axonal water fraction, intra-axonal diffusivity or extra-axonal perpendicular diffusivity hold high clinical value due to their selectivity to key pathophysiological processes like axonal loss^4^, axonal beading^5^ or (de)myelination^4^. Among SM-based formulations, Standard Model Imaging (SMI)^6^ has demonstrated superior specificity when validated against 3D electron microscopy^7^, along with robust and reproducible estimations on standard clinical scanners equipped with low-to-moderate gradient strengths (∼40–80 mT/m).

The recent extension of biophysical modeling to gray matter (GM) marked a paradigm shift, as models originally developed for WM failed to capture the higher soma density, lower myelin content, and consequent greater transmembrane water exchange characteristic of GM tissues^8^. The Neurite Exchange Imaging (NEXI)^8,9^ model, concomitantly introduced also as Standard Model with Exchange (SMEX)^10^, was designed to account for transmembrane water exchange and has been successfully adapted to clinical scanners with 80 mT/m gradients^11^, enabling reliable estimation of intra-/extra-neurite diffusivities, cell-process density, and exchange time between intra-/extra-neurite compartments. These parameters have shown high sensitivity to cortical microstructure, with cell-process density and exchange time capturing the laminar architecture of the hippocampus in postmortem human tissue^12^ and reflecting both neuronal and astrocytic density in rat brains^8^. Furthermore, the transmembrane exchange time, a proxy for membrane permeability, has recently been shown to correlate with Myelin Water Fraction (MWF)^11^, reflecting patterns of cortical myelination in the healthy human brain. In addition, t_ex_ may be sensitive to microstructural changes related to neuronal activity^13^, highlighting its potential for detecting subtle differences across brain regions and functional states.

Magnetization Transfer (MT)–based methods offer an alternative class of contrast mechanisms sensitive to tissue microstructure. Unlike diffusion MRI, which infers tissue organization from the random displacements of water molecules over micrometer-scale distances, MT probes the molecular environment via interactions between free water protons and those bound to macromolecules^14^, making it particularly informative about the macromolecular content such as myelin, but also other lipids and proteins. Accordingly, the macromolecular proton fraction (MPF)^15^, derived from the binary spin-bath model of MT^16^, provides a highly sensitive measure of myelin content^17^, but with limited specificity^18^. A further refinement, inhomogeneous MT (ihMT)^19^, exploits the non-averaged residual dipolar couplings arising preferentially in highly ordered, tightly packed motion-restricted macromolecular structures such as myelin bilayers, and has shown enhanced specificity to myelination^18,20^. IhMT has demonstrated versatility, enabling precise detection of white matter damage^21,22^ and detailed insights into cortical myeloarchitecture^23^. However, like MPF, ihMT is not exclusively specific to myelin; depending on the off-resonance saturation parameters, it can also reflect other macromolecules, including lipid bilayers in glial and neuronal membranes^18^.

Although diffusion and MT contrasts rely on distinct acquisition principles, their biophysical sensitivities partially overlap, resulting in metrics that can capture similar underlying tissue features. At the same time, each modality emphasizes different microstructural aspects, providing complementary insights into the same biological environment from distinct spatio-temporal scales. As such, previous studies have already explored correlations between MT and conventional diffusion metrics derived from DTI^24^, which have relatively low specificity to microstructural features. In the present work, we extend these investigations by systematically assessing associations between metrics derived from advanced diffusion models in both white and gray matter, offering a more detailed characterization of tissue microstructure. We focus on two MT-derived metrics with distinct sensitivity and specificity to myelination^18^—namely, the MPF and the ihMT ratio (ihMTR)—and employ the SMI model in white matter and the SMEX model in gray matter to relate diffusion-derived indices of axonal and neurite composition to these MT measures. This framework allows for a more nuanced understanding of how myelin and cellular microstructure jointly shape MRI contrasts across brain tissue types.

## 2. METHODS

### 2.1. Participants

Ethical approval for the study was obtained from the Ethics Committee of the Canton of Vaud, Switzerland (CER-VD). Written informed consent was obtained from all participants. The cohort comprised 10 healthy adults (8 females, 2 males) with a mean age of 21.3 ± 1.6 years, assessed using an identical protocol.

### 2.1. Data acquisition

All data were collected using a 3T MAGNETOM Prisma MRI system with 80 mT/m gradients (Siemens Healthineers AG, Forchheim, Germany). First, an anatomical reference was acquired using an MP-RAGE sequence (1-mm isotropic resolution, field-of-view 224×240 mm^2^, 256 slices, inversion/repetition time TI/TR = 900/1760 ms). Research dMRI and ihMT acquisition sequences were employed to obtain data for the derivation of the parametric maps of interest. A summary of the relevant acquisition parameters and the corresponding acquisition time for each research sequence is given in **Table 1**.

**Table 1.**
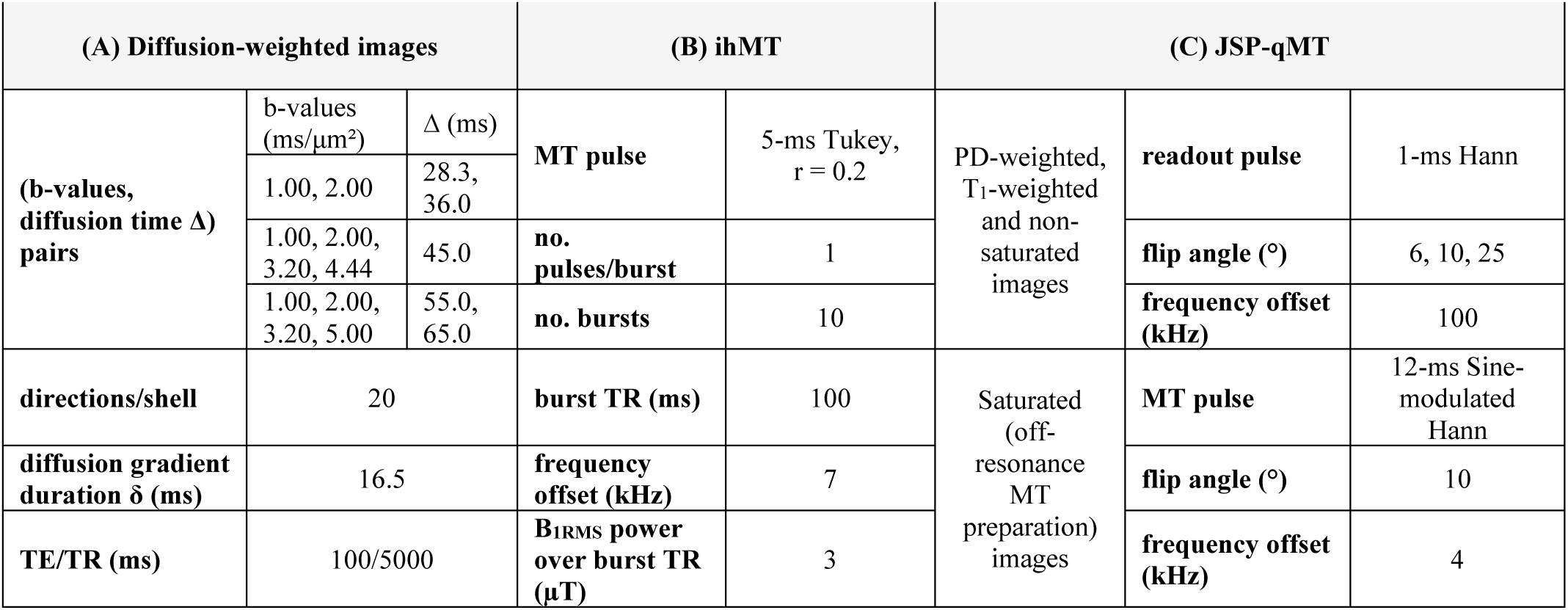
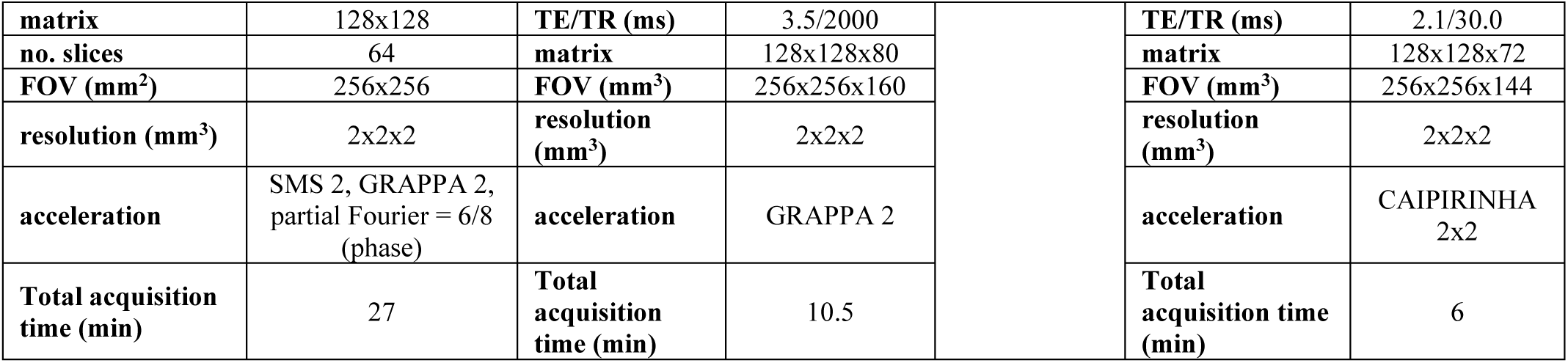
Acquisition parameters for (A) diffusion, (B) ihMT (C) Joint Single-Point – quantitative MT (JSP-qMT) mapping.

#### 2.1.1. Diffusion

A 2D multi-slice Pulsed Gradient Spin Echo sequence with Echo Planar Imaging readout (PGSE-EPI) was used to acquire a multi-shell, multi-diffusion time protocol at 2-mm isotropic resolution, consisting of diffusion-weighted images with varying pairs of b-values and diffusion times (Δ), as previously described^11^. Specifically, for diffusion times Δ = 28.3 ms and 36.0 ms, b-values of 1.00 and 2.00 ms/μm² were collected, for Δ = 45.0 ms, b-values of 1.00, 2.00, 3.20, and 4.44 ms/μm², and for Δ = 55.0 ms and 65.0 ms, b-values of 1.00, 2.00, 3.20, and 5.00 ms/μm², respectively. The gradient duration δ was 16.5 ms. Each shell consisted of 20 directions, and one additional b = 0 (b_0_) image was acquired for each diffusion time. One b = 0 image with reversed EPI phase encode direction was acquired separately for susceptibility distortion correction. Additional readout acquisition variables are indicated in **Table 1A**. The total scan time for the diffusion data was 27 min.

#### 2.1.2. Inhomogeneous Magnetization Transfer

A 3D ihMTRAGE sequence^25^ composed of an ihMT preparation with RApid Gradient Echo (RAGE) readout was employed to acquire four sagittal MT-weighted images and one non-saturated MT_0_ image. The ihMT preparation consisted of 5-ms off-resonance Tukey-shaped pulses (r = 0.2) repeated every 100 ms over a total saturation period of 1 s, with an off-resonance frequency (Δf) of 7 kHz. The MT-weighted images included two single-frequency preparations with either positive (+7 kHz, MT^+^) or negative (−7 kHz, MT^−^) frequency offsets, and two dual-frequency preparations achieved by sine-modulating Tukey-shaped pulses at both offsets (+/–7 kHz, MT^±^ and MT^∓^), applied on-resonance at a B_₁peak_ amplitude scaled by √2 relative to the single-frequency saturation to maintain equivalent power deposition. Readout acquisition variables are detailed in **Table 1B**. The total scan time for ihMT was 10.5 min.

#### 2.1.3. Joint Single Point Quantitative Magnetization Transfer

The imaging protocol consisted of 3D sagittal acquisitions of MT-Spoiled Gradient-Recalled Echo (MT-SPGR) images and variable flip angle (VFA)-SPGR images^26^. One MT-weighted image was acquired using an on-resonance Hann-shaped pulse with a duration of 12.0 ms and a flip angle of 10°, sine-modulated at ±4 kHz to produce simultaneous dual-offset saturation at both +4 kHz and −4 kHz and mitigate for dipolar order effects biasing qMT estimations^19,26,27^. VFA experiments were conducted using the same SPGR sequence, but with the offset frequency of the MT pulse set to 100 kHz in order to maintain a constant RF duty cycle while avoiding creating MT effects. The sequence was repeated three times with flip angles of 6°, 10°, and 25°, to generate one proton density (PD)-weighted, one MT_0_ and one T_1_-weighted image. Common MT-SPGR and VFA-SPGR readout parameters are shown in **Table 1C**. The total acquisition time was 6 min. Additionally, a sagittal interleaved and multislice B_1_^+^ mapping sequence (manufacturer presaturated turbo-FLASH) was used with TE/TR = 2.2/26290.0 ms, BW = 490 Hz/voxel, saturation FA = 80°, readout FA = 8°, in-plane matrix size = 96×96, 60 slices, and reconstructed voxel size = 3×3×3 mm^3^; and acquisition time of 53 s.

### 2.2. Data pre-processing and parametric maps derivation

#### 2.2.1. Diffusion

The multi-shell multi-diffusion time data were pooled together (325 volumes) and underwent the following steps: denoising using the Marchenko-Pastur principal component analysis (MP-PCA) on the magnitude data^28,29^, Gibbs ringing correction^30,31^, magnetic field inhomogeneity distortion correction^32^ and eddy current and motion correction^33^. The high SNR images (b = 0 and b = 1 ms/μm²) were extracted (105 volumes) and denoised separately using MP-PCA to generate unbiased noise maps for Rician mean correction^9,34^.

##### White Matter

The SMI MATLAB (2021a, MathWorks, Natick, Massachusetts) toolbox (https://github.com/NYU-DiffusionMRI/SMI) was used to estimate the SMI parametric maps as previously described^6^. Only the multi-shell diffusion data at Δ= 65 ms was used for the SMI estimation to ensure the long-time limit regime. The calculated maps included: the axonal water fraction (f), the intra-axonal diffusivity (D_a_), the extra-axonal parallel diffusivity (De^||^), the extra-axonal perpendicular diffusivity (D_e_┴) and the axonal dispersion (p_2_).

##### Gray Matter

Prior to estimating the SMEX parametric maps (https://github.com/QuentinUhl/graymatter_swissknife), the data were averaged across all directions for each b-value using the arithmetic mean (powder average) and normalized to the b = 0 volume corresponding to each diffusion time acquisition. The SMEX model accounting for wide gradient pulses and the Rician noise floor^9,11^ was fit to the experimental data using Nonlinear Least Squares (NLS) optimization^11^ with the L-BFGS-B algorithm implemented in *scipy.optimize*^35^ with a tolerance of 10^-14^. The following parametric maps were estimated: cell-process density (f), the intra-/extra-neurite diffusivities (D_i_/D_e_) and the exchange time (t_ex_) between intra-/extra-neurite compartments. The bounds specified for the optimization were [1 - 150] ms for t_ex_, [0.1 - 3.5] µm²/ms for the two diffusivities and [0.1 - 0.9] for the cell-process fraction f. Parameter initialization enforced the condition D_i_ > D_e_, consistent with previous reports^36–38^. The Rician scale σ (which determines the Rician floor as 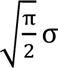) was fixed to the noise standard deviation estimated from the high SNR images during preprocessing.

#### 2.2.2. Inhomogeneous Magnetization Transfer

Images were corrected for Gibbs-ringing artifacts with an isotropic 3D-cosine kernel apodization and motion-corrected^39,40^. The MT-weighted images were combined to calculate the ihMT ratio (ihMTR) maps according to the formula^19^:

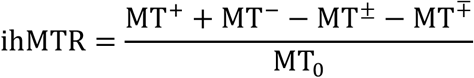

#### 2.2.3. Joint Single-Point quantitative Magnetization Transfer

Each multi-echo SPGR image was combined using a sum-of-squares operation for SNR enhancement. Motion-correction^40^ was performed as follows: VFA- and MT-weighted images were rigidly registered to the VFA-SPGR image acquired at a flip angle of 25°, compensating for intra-scan motion. Quantitative fitting was performed using the joint single-point qMT model, which is unbiased by on-resonance saturation and dipolar order effects^26^, yielding MPF parametric maps.

### 2.4. Analyses

#### 2.4.1. ROI approach

##### 2.4.1.1 Quantification

The ihMTR, MPF and average b_0_ maps for each subject were rigidly registered to their corresponding T_1w_ anatomical reference image. For the b_0_-to-T_1w_ registration, the boundary-based registration method implemented in *epi_reg*^41^ (FMRIB software library; FSL, v.6.0.7) was employed, whereas ihMTR and MPF maps were rigidly registered using the *antsRegistrationSyN.sh* routine^42^ implemented in the Advanced Normalization Tools^42^ (ANTs, v.2.6.2). The co-registered T_1w_, b₀, ihMTR, and MPF maps for each subject were subsequently used to create multivariate templates using the ANTs multivariate template construction workflow (https://github.com/stnava/ANTs/blob/master/Scripts/).

###### White Matter

The MNI template was non-linearly registered to the T_1w_ template generated in this study. The resulting transformations were concatenated with the inverse transformations resulting from the multivariate template construction and the inverse rigid registrations initially applied to the native b_0_, ihMTR and MPF maps, enabling the propagation of the Johns Hopkins University (JHU) white matter atlas^43^ into the native space of each subject and each metric. A white matter mask for each subject was generated using *FAST* on the subject-specific T_1w_ images using FreeSurfer (v7.4.1), slightly eroded, and propagated into the native space of each metric. This mask was then multiplied with the individual JHU segmentation masks to erode them and minimize contamination from cerebrospinal fluid (CSF) or GM for ROI-wise quantifications. Finally, mean values were computed within each JHU ROI for each subject and metric.

###### Gray Matter

Subject-specific native T_1w_ images, denoised and corrected for bias field nonuniform intensity^44^, were processed using FreeSurfer to obtain cortical parcellation^45,46^. The inverse transformations estimated from rigid registration of native b_0_, ihMTR and MPF maps were then applied to the FreeSurfer Desikan-Killiany-Tourville (DKT) segmentation^47^ to generate cortical segmentations for each metric in every subject. Because GM ROIs are relatively small, median values were computed within each ROI to ensure robustness against outliers.

##### 2.4.1.2 Statistical analyses

Average values across the ten subjects were computed for each metric, and Pearson’s correlation coefficients were calculated for pairs of ihMTR and MPF with SMI and SMEX parameters using the Python *pingouin* library (v5.5)^50^. False discovery rate (FDR) correction was applied to account for multiple comparisons in the SMI and SMEX correlations.

#### 2.4.2. Surface analyses

The white matter and pial surfaces were extracted from the FreeSurfer analysis of the T_1w_ template generated in this study to create an inflated brain. All parametric maps derived from SMI and SMEX, as well as ihMTR and MPF, were propagated into the space of the multivariate templates, ensuring alignment with T_1w_ template. For visualization of regional distributions, each metric was first averaged across all subjects and then projected onto the cortical surfaces.

## 3. RESULTS

### 3.1 Metrics values from white matter and gray matter regions

**Figure 1** presents axial slices of the b₀, ihMTR, and MPF templates averaged across the 10 participants included in this study, alongside the T_1_-weighted anatomical template with the corresponding WM and GM segmentations overlaid.

**Figure 1.**
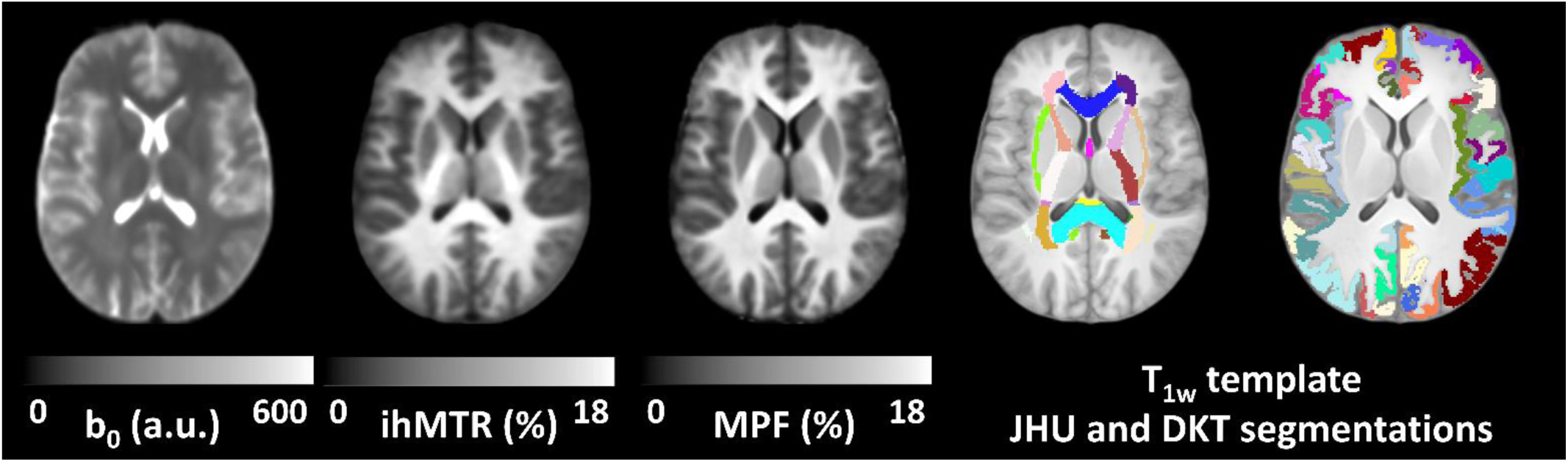
Multivariate templates generated from the b_0_, ihMTR, MPF, and T_1w_ images of the 10 subjects included in this study. The T_1w_ and MNI templates were non-linearly co-registered, enabling the propagation of the JHU white matter atlas to the study-specific multivariate template space, and subsequently into each subject’s native space for all metrics. The native T_1w_ images of each subject were processed with Freesurfer to generate DKT segmentations of the cortical ribbon. The DKT segmentation of the T_1w_ template is illustrated here as an example.

Although ihMTR and MPF exhibit broadly similar contrasts, several regional distinctions emerge as summarized in **Table S1** and **Table S2** in Supplementary Information. For example, in WM (**Table S1**), ihMTR values are higher in the posterior limb of the internal capsule and the corticospinal tract, whereas MPF shows greater values in the genu and splenium of the corpus callosum. Notably, ihMTR demonstrates a gradual increase along the corpus callosum—from the genu through the body to the splenium. These regional differences are further illustrated in **Figure 2** on average ihMTR and MPF maps of the 10 subjects. Overall, the WM distribution of ihMTR is slightly shifted toward higher values compared with MPF, as illustrated by the scatter plots in **Figure 3**, with both its minimum and maximum values exceeding those of MPF. In GM (**Table S2**), ihMTR and MPF also differ regionally and in their overall range of values, with ihMTR starting at lower values but reaching higher maxima (**Figure 3**). Cortical regions such as cuneus and the superior temporal gyrus have high ihMTR, while precentral gyrus and the superior frontal gyrus present high MPF.

**Figure 2.**
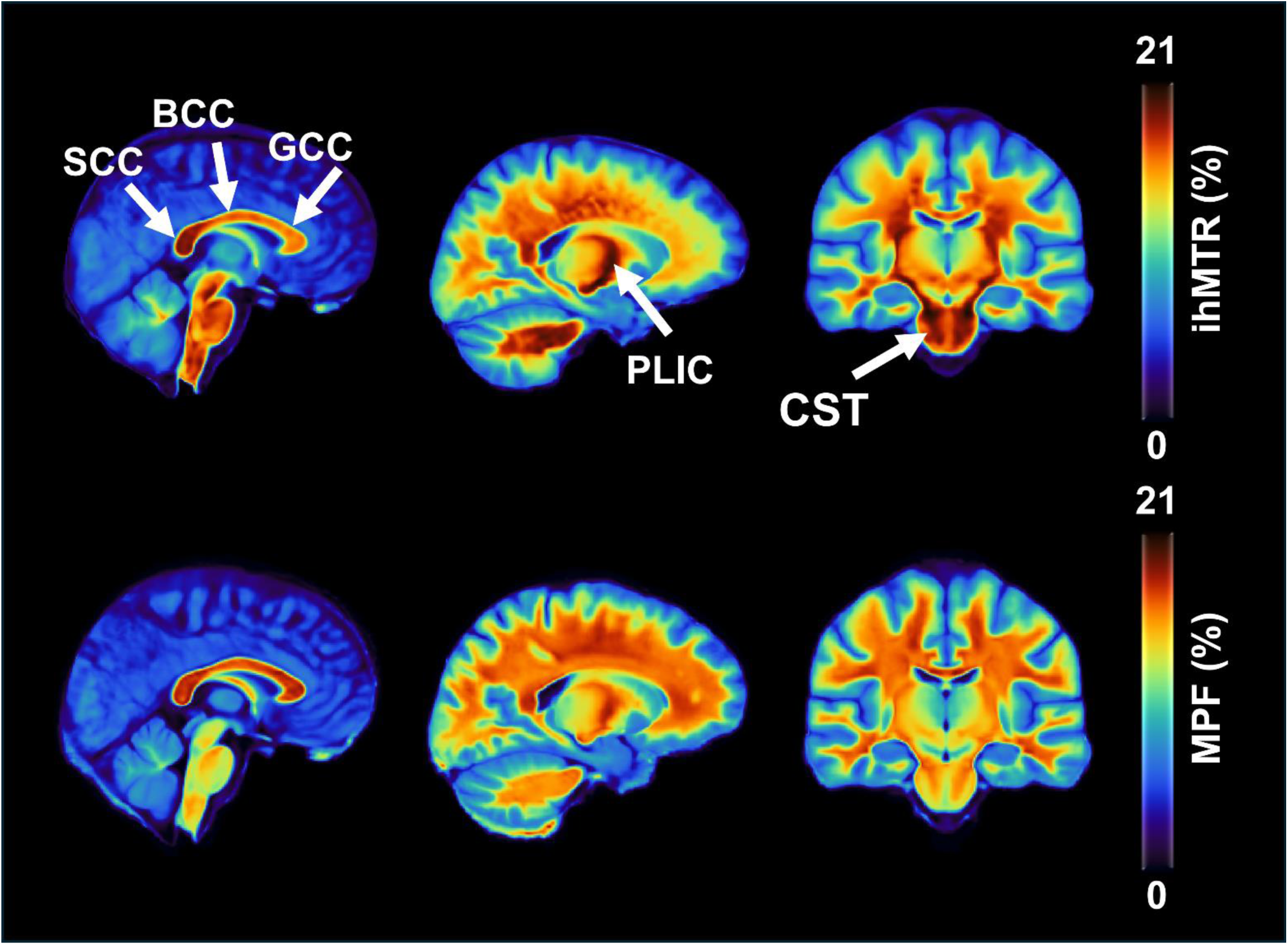
Example of ihMTR versus MPF–specific differences in WM brain regions. An increasing anterior-to-posterior gradient of ihMTR is visible along the corpus callosum, with progressively higher values in the genu (GCC), body (BCC), and splenium (SCC), in contrast to the high–low–high pattern observed for MPF. The corticospinal tract (CST) and the posterior limb of the internal capsule (PLIC) showed ihMTR values comparable to those in the SCC, whereas MPF values in these same brain structures were lower than in SCC and GCC.

**Figure 3.**
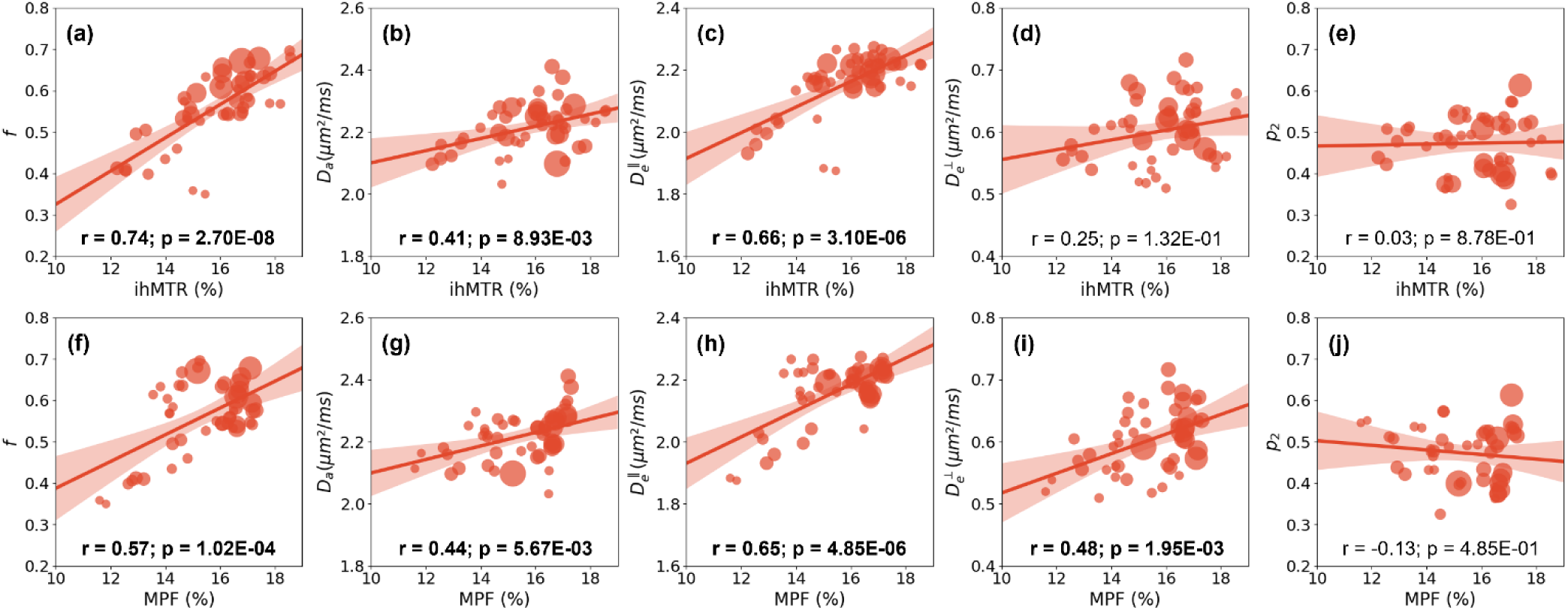
Scatter plots showing the associations between ihMTR and MPF with the SMI-derived metrics. f – axonal water fraction, D_a_ – axonal water diffusivity, D_e_^||^ –extracellular parallel diffusivity, D_e_^┴^ –extracellular perpendicular diffusivity, and p_2_ – axonal dispersion. Linear regression lines with 95% confidence intervals are displayed for visual guidance. Pearson’s correlation coefficients (r) and corresponding p-values (p) FDR-corrected for multiple comparisons are indicated in each plot; values shown in **bold** denote significant correlations after FDR correction.

To summarize, across WM ROIs, ihMTR ranged from 12.2–18.6% and MPF from 11.6–17.3%; in GM ROIs, ihMTR ranged from 5.9–8.6% and MPF from 6.3–8.4%. For SMI, parameter ranges were as follows: f: 0.35–0.70, Dₐ: 2.03–2.41 µm²/ms, Dₑ^||:^ 1.87–2.27 µm²/ms, Dₑ^⊥^: 0.51–0.72 µm²/ms, and p₂: 0.33–0.61. For SMEX, the corresponding ranges were: f: 0.32–0.51, tₑₓ: 16.7–40.5 ms, Dᵢ: 2.10–3.09 µm²/ms, and Dₑ: 0.90–1.59 µm²/ms.

### 3.2 Microstructural correlates in white matter ROIs

**Figure 3** presents the scatter plots along with the linear regression line, Pearson’s correlation coefficients (r) and corresponding FDR corrected p-values (p) for associations between pairs of ihMTR and MPF with the five SMI metrics. Each point in the scatter plots represents the average across 10 subjects, where the average within each ROI was calculated. The size of the points reflects the relative size of the ROIs, with smaller points corresponding to regions such as the tapetum, which also deviated most from the linear regression line. In WM ROIs, ihMTR showed a very strong correlation with the axonal water fraction f (r = 0.74, p < 0.001), a strong correlation with the extra-axonal parallel diffusivity D_e_^||^ (r = 0.66, p < 0.001), and a moderate correlation with the intra-axonal diffusivity D_a_ (r = 0.41, p < 0.01). Correlations with the extra-axonal perpendicular diffusivity D_e_^⟂^ were weak (r = 0.25, p = 0.13) and not statistically significant, while associations with the axonal dispersion p_2_ were negligible and non-significant. When assessed against MPF, all correlations were statistically significant, except for the axonal dispersion p_2_. Specifically, correlations were strong for the axonal water fraction f (r = 0.57, p < 0.001) and the extra-axonal parallel diffusivity D_e_^||^ (r = 0.65, p < 0.001), and moderate for diffusivities D_a_ (r = 0.44, p < 0.01) and D_e_^⟂^ (r = 0.48, p < 0.01). The confidence intervals and p-values before and after correction for multiple comparisons are displayed in **Table S3** and **Table S4** (Supplementary Information) for associations with ihMTR and MPF, respectively.

**Figure 4** shows axial slices of the group-average maps for SMI, ihMTR, and MPF metrics across ten subjects, with maps of WM ROIs (JHU atlas) overlaid in color for clearer visualization. The spatial patterns reveal clear regional differences between ihMTR and MPF, as well as a stronger spatial correspondence between f and ihMTR than between f and MPF.

**Figure 4.**
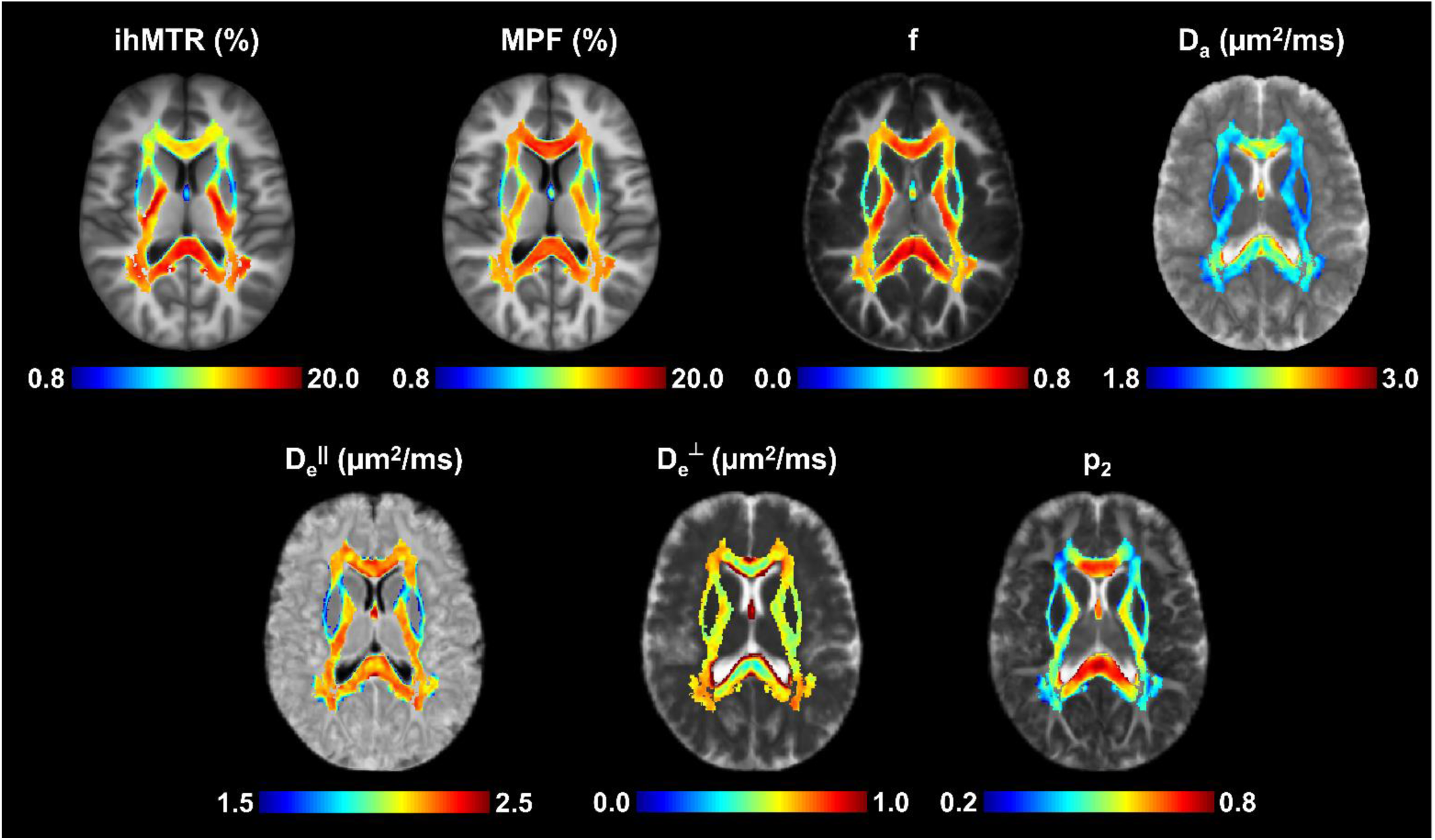
Axial slices from the 3D voxel-wise group-average maps of SMI, ihMTR, and MPF metrics derived from the ten participants. The extent of the colormaps is limited to the ensemble of JHU ROIs to limit the dynamic range of the metrics and enhance visualization. The maps reveal distinct spatial patterns for each metric, with ihMTR more closely mirroring the distribution of the axonal water fraction f.

### 3.3 Microstructural correlates in the gray matter cortical ribbon

**Figure 5** presents the scatter plots along with the linear regression line, Pearson’s correlation coefficients (r) and corresponding FDR corrected p-values (p) for associations between pairs of ihMTR and MPF with the four SMEX metrics. Each point in the scatter plots represents the average across 10 subjects, where the median within each ROI was calculated for robustness against outliers. Again, the size of the points reflects the relative size of the ROIs, with smaller points corresponding to regions such as the caudal anterior cingulate, entorhinal cortex, and pericalcarine cortex, which also tended to deviate the most from the linear regression line. Interestingly, the exchange time t_ex_ exhibited a strong significant correlation with ihMTR (r = 0.57, p < 0.001), while the cell-process fraction f showed a significant moderate-to-strong correlation with MPF (r = 0.49, p < 0.001). Associations between ihMTR and f, and MPF and t_ex_ were not statistically significant. Intra-neurite diffusivity D_i_ and extra-neurite diffusivity D_e_ demonstrated weak correlations with MPF (r = 0.27, p = 0.053) and ihMTR (r = 0.27, p = 0.051) respectively, though they did not survive FDR correction for multiple comparisons. Correlations between D_e_ and MPF, as well as between D_i_ and ihMTR, were negligible and not statistically significant. The confidence intervals and p-values before and after FDR correction for multiple comparisons are displayed in **Table S4** (Supplementary Information).

**Figure 5.**
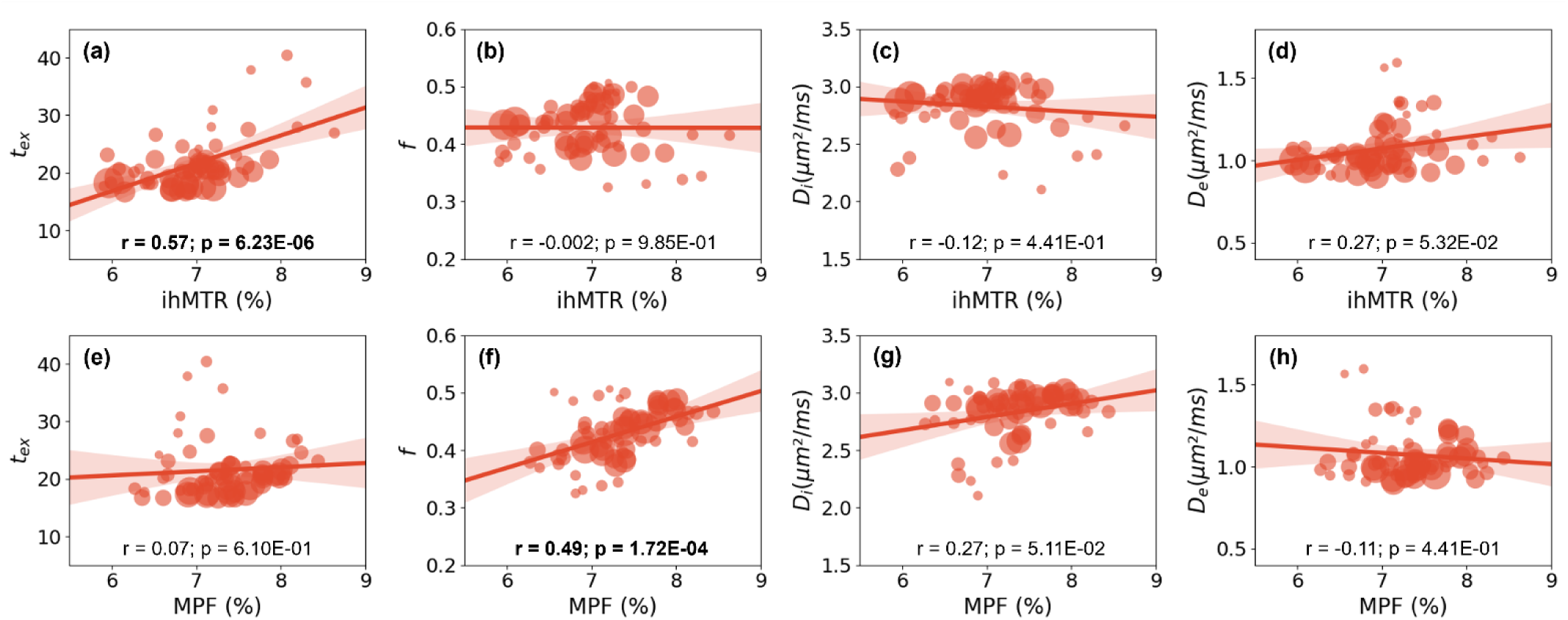
Scatter plots showing the relationships between ihMTR and MPF with the SMEX-derived metrics: t_ex_ – exchange time, f – cell-process fraction, D_i_ – intra-neurite diffusivity, and D_e_ – extra-neurite diffusivity. Linear regression lines with 95% confidence intervals are displayed for visual guidance. Pearson’s correlation coefficients (r) and corresponding p-values (p) FDR-corrected for multiple comparisons are indicated in each plot; values shown in **bold** denote significant correlations after FDR correction.

**Figure 6** shows cortical surface projections highlighting highly myelinated regions around the central sulcus and occipital lobe, which exhibit longer tₑₓ and higher f, ihMTR, and MPF values. Interestingly Di was also high, while De was low in the same region. Discrepancies observed in other regions, such as the temporal cortex, may reflect regional differences in microstructural composition or measurement sensitivity.

**Figure 6.**
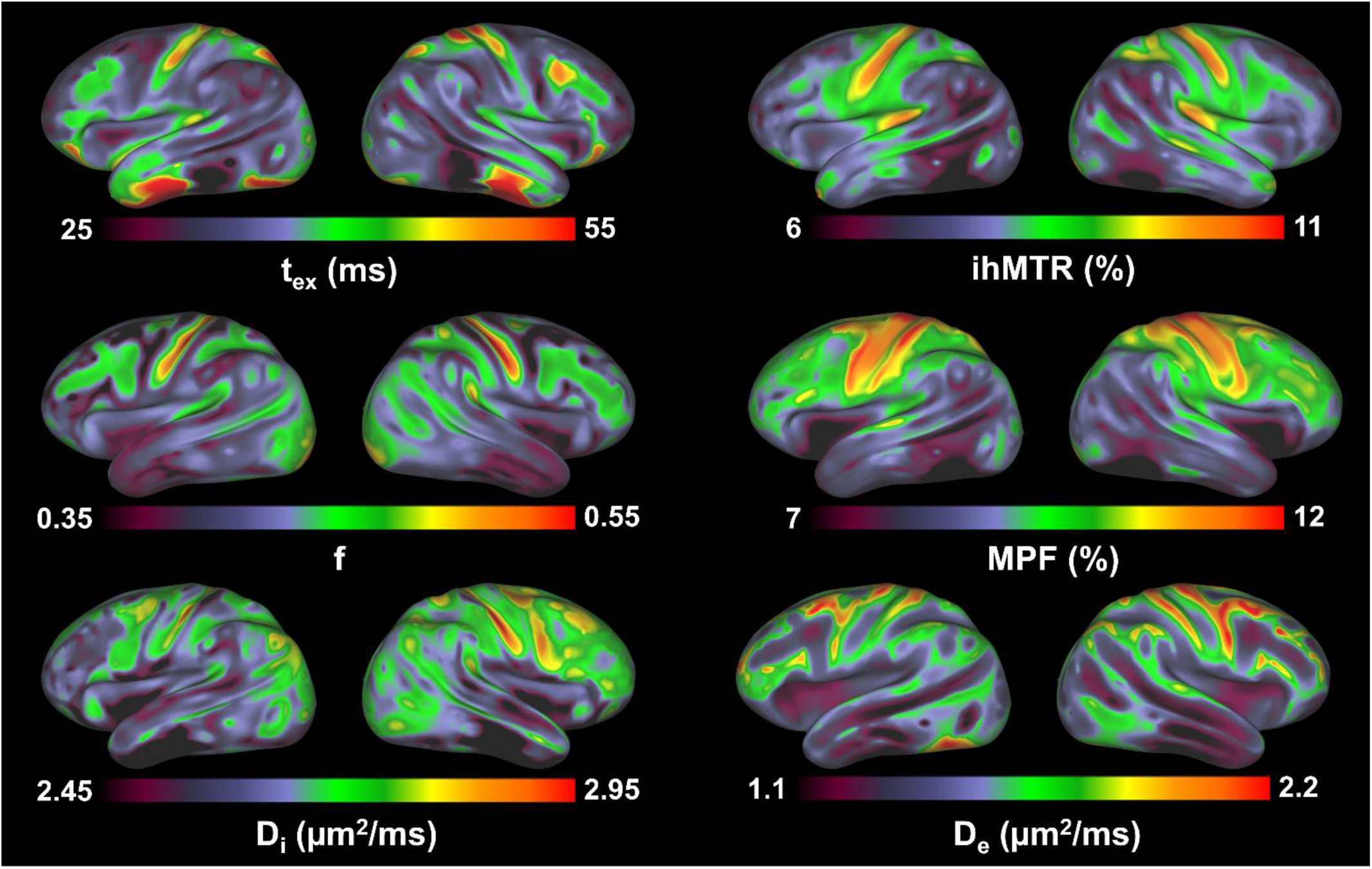
Cortical surface projections of SMEX-derived parametric maps, MPF, and ihMTR. The primary motor cortex stands out due to its systematic variations across metrics, with longer tₑₓ, higher f, D_i_, ihMTR, and MPF, and lower D_e_. Discrepancies in other brain regions (occipital and temporal lobes) likely reflect the distinct underlying sensitivity of each metric to the local microstructural composition.

## 4. DISCUSSION

In this study, we investigated the complementarity between metrics derived from advanced biophysical models of diffusion MRI and two MT-based techniques, qMT and ihMT. White and gray matter brain regions were analyzed separately because of their distinct microstructural organization and biophysical properties. In white matter, five SMI-derived metrics—capturing intra- and extra-axonal (parallel and perpendicular) diffusivities, axonal dispersion, and axonal water fraction—were correlated with MPF and ihMTR. In gray matter, we assessed correlations between MPF and ihMTR with four SMEX-derived metrics: intra- and extra-neurite diffusivities, cell-process density, and exchange time between the intra-/extra-neurite compartments.

### 4.1 Microstructural correlates in white matter fiber tracts

The SMI parametric maps in this study are consistent with previous healthy human brain reports^6^. Small differences in the estimates may reflect dissimilitude in study populations, such as age^49^, or methodological dissimilitude, particularly in the diffusion weighting and readout parameters like the echo time, previously shown to affect SMI estimates^6^. Likewise, ihMTR and MPF maps obtained across the investigated WM tracts are consistent with previous studies in the healthy human brain^25,26^. Despite their similar WM/GM contrast and shared biophysical underpinnings, ihMT and MPF exhibited several notable differences. The highest ihMTR values were observed in myelin-rich brain structures, such as the corticospinal tract, the posterior limb of internal capsules and the corpus callosum. Interestingly, an ihMTR gradient was observed in the longitudinal axis of the corpus callosum, with increasing values from genu to body to splenium, consistent with previous histological studies of myelin density^50^ and MRI studies measuring MWF^51^ or the paramagnetic myelin signals in quantitative susceptibility imaging^52^. Notably, ihMTR was higher in the corticospinal tract than in the corpus callosum, possibly reflecting a greater myelination of individual axons. However, some of the ihMTR differences across ROIs may partly reflect orientation effects in addition to genuine variations in myelin content. Because ihMT contrast depends on fiber orientation relative to the main magnetic field B_0_^24,53^, fibers that are oriented very differently—such as the corpus callosum and the corticospinal tract, which are approximately perpendicular to each other—may exhibit contrast differences that are not solely attributable to myelin. On the other hand, MPF displayed a high-low-high pattern in the longitudinal axis of the corpus callosum as well as higher overall values than in the corticospinal tract or internal capsules. MPF patterns in this study align with prior findings^26^ and differ markedly from the corresponding ihMT profiles.

Correlations between SMI parameters with ihMTR and MPF revealed significant associations across all metrics, with the exception of axonal dispersion p₂ with both ihMTR and MPF, and extra-axonal perpendicular diffusivity D_e_^⊥^ with ihMTR. The positive correlations of axonal water fraction f against ihMTR and MPF likely reflect that brain regions with higher axonal water fraction also contain more myelin, thereby enhancing both ihMTR and MPF^17^. This interpretation is supported by prior electron-microscopy studies showing that axonal water fraction closely tracks not only the total number of axons but also the subset of myelinated axons and myelin volume fraction^4^. Remarkably, the stronger correlation observed in this study between axonal water fraction with ihMTR reflects the higher specificity of ihMT to myelin content as compared to MPF^18^. Furthermore, a positive association between MPF and histology-derived axonal density was previously reported^54^, although weaker than that observed with myelin. This may explain the dissimilar patterns between ihMTR and MPF observed in particular in the corpus callosum and may point to a more complex relationship between the measured MPF and the underlying tissue architecture. In addition to myelin content, MPF is likely modulated by a combination of axon density, size and spacing, all impacting axonal packing. For instance, a voxel containing a few large axons may have the same axonal water fraction as one with many small axons, yet the overall macromolecular fraction would differ due to variations in membrane surface area contributing to MT effects.

As for water compartment-specific diffusivities, both the intra-axonal diffusivity D_a_ and extra-axonal parallel diffusivity D_e_^||^—capturing complementary intra- and extra-axonal features of axonal beading, undulation and orientation coherence^5,55,56^—exhibited similar positive correlations with ihMTR and MPF. Given that higher parallel diffusivities are measured for coherently packed, straight axons, a morphology promoted by strong myelination, it is not unexpected that both diffusivity metrics exhibited comparable associations with ihMTR (through myelin and fiber orientation) and MPF (through myelin and most likely fiber packing). More surprisingly though, the extra-axonal perpendicular diffusivity D_e_^⊥^—a parameter known to reflect myelin content changes in pathological conditions^4^—showed positive correlation with MPF. The correlation between D_e_^⊥^ and ihMTR was also positive, but weak and non-significant. Since a higher degree of myelination restricts radial water diffusivity, D_e_^⊥^ should decrease as myelin content increases; accordingly, one would expect negative, rather than positive, associations between D_e_^⊥^ and MPF/ihMTR. However, in healthy tissue, variations in D_e_^⊥^ across fiber tracts may be less indicative of myelin content and more reflective of fiber coherence and packing, as suggested by the significant correlations of D_e_^⊥^ with extra-axonal parallel diffusivity (D_e_^||^, r = 0.43, p < 0.01) and intra-axonal diffusivity (D_a_, r = 0.47, p < 0.01). This likely explains the significant correlation between D_e_^⊥^ and MPF only, further reinforcing the idea that MPF captures aspects of fiber packing in addition to myelin content. Future studies combining MPF measurements with detailed axonal properties derived from electron microscopy will be essential to clarify the precise relationship between MPF and axonal architecture.

Remarkably, neither ihMTR nor MPF correlated with the axonal dispersion p_2_. Although numerical simulations have shown that ihMTR can be influenced by axonal dispersion^57^, this effect is likely modest under in-vivo conditions, where substantial within-voxel heterogeneity in fiber architecture tends to average out dispersion-related contributions. As a result, stronger determinants—such as myelin content and fiber-orientation–dependent MT effects—dominate the measured ihMTR. A similar reasoning applies to MPF: any potential influence of axonal dispersion is probably small and effectively overshadowed by MPF’s much greater sensitivity to myelin content and axonal packing.

### 4.2 Microstructural correlates in the gray matter cortical ribbon

The SMEX parameters obtained in this study, along with their cortical surface projections, are consistent with previous reports in the healthy human brain^9,11,58,59^. SMEX cortical surface projections exhibited consistent variations in the primary motor cortex, with the exchange time tₑₓ, the cell-process fraction f, and intra-neurite diffusivity Dᵢ showing high values, and the extra-neurite diffusivity Dₑ showing low values; a pattern paralleled by increased ihMTR and MPF. However, differences between all the metrics are apparent brain-wide, likely reflecting their distinct sensitivities to local microstructural composition.

Regarding correlations between SMEX parameters and ihMTR/MPF, only two remained significant after FDR correction: a positive association between tₑₓ and ihMTR, and between MPF and Dᵢ. The correlation between tₑₓ and ihMTR aligns with previous reports of strong associations between tₑₓ and T₂-derived MWF^11^, supporting a link between tₑₓ and myelination. Our results are therefore consistent with the notion that higher myelination (i.e., higher ihMTR) is associated with reduced membrane permeability and longer exchange times. Nevertheless, the association between t_ex_ and ihMTR might not be solely driven by myelination. Although the ihMTR acquisition used in this study was previously applied for cortical myelination mapping^23^ and showed good correspondence with other myelin-sensitive measures^60,61^, some contribution from glial cells^18,21^ cannot be excluded given their higher density in gray matter relative to white matter. Conversely, tₑₓ reflects the overall exchange dynamics between the intra- and extra-neurite compartments and therefore lacks cellular specificity, with previous studies illustrating its sensitivity to astrocytic density^8^. Discrepancies between t_ex_ and ihMTR, in cortical surface projections (e.g., sensory area or occipital and temporal lobes) might arise from distinct sensitivities of these metrics to other factors. Specifically, t_ex_ is influenced by dendritic spine complexity^62^, while ihMTR can be affected by T_1_ relaxation during the off-resonance saturation module and B_1_^+^ inhomogeneity introduced by the excitation pulses^23^.

Less expected, the lack of correlation between MPF and tₑₓ may reflect a shift in the dominant microstructural feature influencing MPF in gray matter. Compared to white matter, correlations between MPF and histological myelin markers have been shown to be weaker, while associations with cellular density became stronger in gray matter^54^. The greater contribution of bound protons from glial cells and their processes may therefore explain the positive correlation found in this study between MPF and the cell-process fraction *f*.

### 4.3 ihMTR vs. MPF

The weak-to-moderate correlations (**Figure S1,** Supplementary Information) between ihMTR and MPF observed in WM (r = 0.36, p = 0.02) and GM (r = 0.32, p = 0.02) may appear counterintuitive, given their similar WM/GM contrast and the fact that both metrics arise from interactions between free water and macromolecular protons and are sensitive to tissue macromolecular composition. However, MPF reflects the overall fraction of bound protons relative to the total proton pool and is influenced by multiple microstructural factors, including myelin density, total cellular density, local water content, and probably, axonal packing. On the other hand, ihMTR benefits from additional specificity to dipolar order effects arising from restricted motion and spatial orientation of macromolecular dipolar-coupled protons^19^, providing contrast that depends strongly on microstructural organization in addition to the macromolecular content. The off-resonance saturation scheme in ihMT allows further modulation of macromolecular contributions based on their distinct dipolar order signatures^18,63^. In brain tissues, the existence of at least two dipolar order reservoirs with characteristic dipolar order relaxation time T_1D_ was shown^64^, with the longer component, in the order of tens of milliseconds, being strongly related to myelin^18^. Various acquisition strategies—also referred to as T_1D_-filtering—have been developed to selectively isolate the long T_1D_ components^63^, but increased specificity comes at the cost of reduced sensitivity^18^. In this study, we therefore adopted an ihMT sequence design that strikes a balance between specificity and sensitivity, particularly since ihMT values in gray matter are at least twofold lower than in white matter. However, this approach inevitably preserves some non-myelin contributions with short T_1D_ (in the order of hundreds of microseconds^64^), likely arising from glial cell membranes^18,21^. In gray matter, where glial cell density is higher than in white matter, these non-myelin contributions may be more prominent to the ihMT signal. Despite this, the specificity of ihMTR to myelination, even with lower performance T_1D_ filtering strategies, such as used in this work, was shown to be superior to that of MPF^18^. This is also consistent with previous studies comparing ihMTR or MPF to other myelin sensitive metrics such as MWF. Strong significant correlations between MWF and ihMTR have been reported in the healthy human brain^24^, whereas correlations with MPF in healthy rodent brains were minimal and became significant only in pathological tissue, where myelin content varies more substantially and other microstructural components may also be altered^65^. The lower specificity of MPF to myelination, as well as the distinct microstructural contributions to ihMTR and MPF might therefore explain the modest correlation between the two metrics.

### 4.4. Limitations

Several limitations of this study should be noted. Because the spatial resolution was only 2×2×2 mm³, voxel values in both WM and GM were susceptible to partial volume contamination from neighboring CSF as well as adjacent GM or WM voxels. In WM, where structures are thicker, masks could be safely eroded without substantial voxel loss, thereby reducing these effects. Consistent with this, **Figure S2** shows generally low coefficients of variation across WM regions for all metrics. However, mask erosion was not feasible in the relatively thin cortical ribbon. Partial volume effects were particularly pronounced for the exchange time t_ex_: contamination from WM led to overestimated values—sometimes exceeding 100 ms, more than five times larger than the median cortical t_ex_ value (**Table S2**). In contrast, other SMEX parameters as well as ihMTR and MPF were less affected, with partial volume introducing only comparatively small shifts. This is also reflected in **Figure S3**, which shows the markedly higher coefficient of variation for t_ex_ relative to the other metrics. To reduce sensitivity to extreme values within each ROI, median metrics were computed. Although this approach does not fully eliminate bias, the resulting values were consistent with those reported by other groups using similar resolution^9,11,58^. Furthermore, recent work found no correlation between t_ex_ and cortical thickness at the same spatial resolution^11^, supporting the conclusion that the spatial patterns are driven by the intrinsic biological sensitivity of the method rather than by partial volume effects. Future studies with higher spatial resolution would however facilitate more accurate computation and estimation of the various metrics and a better assessment of their association within the cortical ribbon.

While the absence of ground-truth measurements (postmortem histology) limits the conclusions of this study, our results highlight several important directions for future work. Subsequent studies should aim to disentangle the relative contributions of myelin versus the combined effects of axonal density, diameter, and spacing, particularly in relation to MPF. The influence of various glial cells populations on both ihMTR and MPF also warrants further investigations, along with the effect of surface-to-volume ratio on diffusion-and MT-derived metrics. While WM biophysical models have benefited from various comparison studies against electron microscopy^4,7^, the SMEX parameters so far undergone only qualitative comparisons with histology^8^. Further work is therefore needed to establish the full sensitivity and specificity of the exchange time, the cell-process fraction, and the intra-/extra-neurite diffusivities.

Lastly, our study is subject to biases arising from T_1_ relaxation during the off-resonance saturation module and from B₁⁺ inhomogeneity introduced by the excitation pulses. We chose not to apply the two-step T₁/B₁⁺ correction applied in previous studies^18,23^ because of the multi-compartmental nature of T_1_^66–68^ and its complex interaction with MT measures^69^. Future methods that explicitly model this complexity will be needed to achieve more accurate ihMT quantification in the brain without the T_1_/B₁⁺-related biases.

## 5. CONCLUSION

This work represents the first systematic investigation of the associations between parameters derived from advanced biophysical diffusion models and MT–based measures (ihMTR and MPF) in the healthy human white and gray matter. We provide reference values for SMI and SMEX/NEXI parameters, ihMTR, and MPF across both white matter tracts and the cortical ribbon, along with their correlations. This cross-validation highlights the agreement but also the complementarity of diffusion- and MT-derived metrics, demonstrating that while each parameter is sensitive to multiple microstructural features, their interrelationships have the potential to boost specificity and clarify the baseline contributions of myelin, axonal/cellular density, and other tissue properties. This is especially important in pathological conditions, where the primary disease process is likely to drive observed changes. Establishing baseline correlations thus provides a critical reference for interpreting disease-related alterations and for more accurately identifying the underlying pathological mechanisms. Overall, this study provides a comprehensive reference framework for advanced diffusion and MT metrics in the healthy human brain, supporting their use in future investigations of brain microstructure and pathology.

## Acknowledgements

This work was supported by the Swiss National Science Foundation under Eccellenza grant PCEFP2_194260, and took place at the CIBM Center for Biomedical Imaging, a Swiss research center founded and supported by Lausanne University Hospital (CHUV), the University of Lausanne (UNIL), the Swiss Federal Institute of Technology (EPFL), the University of Geneva (UNIGE) and Geneva University Hospital (HUG). The research ihMTRAGE and vibeMT sequences were provided by the CRMBM laboratory (Aix Marseille University, CNRS) and are available on Siemens’ C2P platform. The authors thank Olivier Girard, Lucas Soustelle, Guillaume Duhamel and Thomas Troalen for sharing the sequences. The authors also wish to thank Jean-Baptiste Ledoux for security training of AH on the 3T scanner at CHUV.

