## Supplementary Information for "Microstructural correlates of white and gray matter in the healthy human brain: comparative analysis of diffusion biophysical models, inhomogeneous magnetization transfer, and macromolecular proton fraction"

**Table S1: Average and standard deviation values of the investigated metrics calculated across 10 subjects in representative WM tracts. Abbreviations:** GCC, genu of corpus callosum; BCC, body of corpus callosum; SCC, splenium of corpus callosum; ALIC-L, anterior limb of internal capsule left; ALIC-R, anterior limb of internal capsule right; PLIC-L, posterior limb of internal capsule left; PLIC-R, posterior limb of internal capsule right; RLIC-L, retrolenticular part of internal capsule left; RLIC-R, retrolenticular part of internal capsule right; CGC-L, cingulum (cingulate gyrus) left; CGC-R, cingulum (cingulate gyrus) right; SLF-L, superior longitudinal fasciculus left; SLF-R, superior longitudinal fasciculus right; UF-L, uncinate fasciculus left; UF-R, uncinate fasciculus right; CST-L, corticospinal tract left; CST-R, corticospinal tract right.

| WM tracts | ihMTR (%) | MPF (%) | f | $D_a$ (μm²/ms) | $D_e^{\parallel}$ (μm²/ms) | $D_e^{\perp}$ (μm²/ms) | $p_2$ |
|---|---|---|---|---|---|---|---|
| **GCC** | 15.1 ± 0.7 | 17.2 ± 0.7 | 0.59 ± 0.03 | 2.28 ± 0.03 | 2.22 ± 0.03 | 0.59 ± 0.04 | 0.54 ± 0.02 |
| **BCC** | 16.0 ± 0.5 | 16.6 ± 0.6 | 0.61 ± 0.03 | 2.25 ± 0.04 | 2.21 ± 0.03 | 0.62 ± 0.03 | 0.51 ± 0.04 |
| **SCC** | 17.4 ± 0.5 | 17.1 ± 0.5 | 0.68 ± 0.02 | 2.29 ± 0.04 | 2.24 ± 0.03 | 0.57 ± 0.03 | 0.61 ± 0.02 |
| **ALIC-L** | 12.9 ± 0.6 | 14.3 ± 0.6 | 0.50 ± 0.02 | 2.12 ± 0.03 | 1.99 ± 0.04 | 0.56 ± 0.03 | 0.48 ± 0.01 |
| **ALIC-R** | 13.3 ± 0.4 | 14.6 ± 0.7 | 0.50 ± 0.02 | 2.16 ± 0.03 | 2.04 ± 0.04 | 0.54 ± 0.03 | 0.50 ± 0.01 |
| **PLIC-L** | 17.6 ± 0.6 | 16.1 ± 0.6 | 0.63 ± 0.02 | 2.15 ± 0.03 | 2.22 ± 0.03 | 0.56 ± 0.04 | 0.52 ± 0.01 |
| **PLIC-R** | 17.8 ± 0.7 | 16.2 ± 0.6 | 0.64 ± 0.01 | 2.15 ± 0.04 | 2.22 ± 0.03 | 0.56 ± 0.02 | 0.52 ± 0.01 |
| **RLIC-L** | 16.3 ± 0.6 | 16.3 ± 0.5 | 0.54 ± 0.03 | 2.22 ± 0.02 | 2.24 ± 0.03 | 0.57 ± 0.03 | 0.53 ± 0.02 |
| **RLIC-R** | 16.8 ± 0.6 | 16.4 ± 0.5 | 0.56 ± 0.02 | 2.24 ± 0.02 | 2.27 ± 0.03 | 0.63 ± 0.03 | 0.51 ± 0.02 |
| **CGC-L** | 14.7 ± 0.7 | 16.6 ± 0.7 | 0.58 ± 0.03 | 2.28 ± 0.05 | 2.17 ± 0.04 | 0.62 ± 0.05 | 0.49 ± 0.03 |
| **CGC-R** | 14.9 ± 0.6 | 16.3 ± 0.7 | 0.56 ± 0.04 | 2.25 ± 0.06 | 2.18 ± 0.07 | 0.65 ± 0.03 | 0.47 ± 0.03 |
| **SLF-L** | 16.1 ± 0.5 | 16.7 ± 0.5 | 0.64 ± 0.03 | 2.27 ± 0.04 | 2.14 ± 0.04 | 0.62 ± 0.04 | 0.41 ± 0.01 |
| **SLF-R** | 16.1 ± 0.4 | 16.8 ± 0.7 | 0.66 ± 0.02 | 2.28 ± 0.04 | 2.15 ± 0.02 | 0.64 ± 0.03 | 0.43 ± 0.01 |
| **UF-L** | 12.6 ± 1.4 | 12.8 ± 0.8 | 0.41 ± 0.04 | 2.16 ± 0.04 | 2.01 ± 0.07 | 0.57 ± 0.04 | 0.51 ± 0.02 |
| **UF-R** | 13.4 ± 1.7 | 12.6 ± 0.6 | 0.40 ± 0.03 | 2.18 ± 0.03 | 2.03 ± 0.05 | 0.60 ± 0.05 | 0.51 ± 0.03 |
| **CST-L** | 18.5 ± 0.9 | 15.3 ± 0.7 | 0.70 ± 0.02 | 2.27 ± 0.08 | 2.22 ± 0.05 | 0.63 ± 0.06 | 0.40 ± 0.02 |
| **CST-R** | 18.6 ± 1.1 | 15.2 ± 0.5 | 0.68 ± 0.01 | 2.27 ± 0.06 | 2.22 ± 0.03 | 0.66 ± 0.05 | 0.40 ± 0.02 |
| **All tracts** | 15.8 ± 1.6 | 15.3 ± 1.5 | 0.56 ± 0.09 | 2.21 ± 0.08 | 2.16 ± 0.10 | 0.60 ± 0.05 | 0.47 ± 0.07 |

**Table S2: Average and standard deviation values of the investigated metrics calculated across 10 subjects in representative GM cortical regions. Abbreviations:** PC-L, pericalcarine cortex left; PC-R, pericalcarine cortex right; PreCG-L, precentral gyrus left; PreCG-R, precentral gyrus right; PCG-L, postcentral gyrus left; PCG-R, postcentral gyrus right; CUN-L, cuneus left; CUN-R, cuneus right; STG-L, superior temporal gyrus left; STG-R, superior temporal gyrus right; MTG-L, middle temporal gyrus left; MTG-R, middle temporal gyrus right; IPL-L, inferior parietal lobule left; IPL-R inferior parietal lobule right; SFG-L, superior frontal gyrus left; SFG-R, superior frontal gyrus right.

| GM ROI | ihMTR (%) | MPF (%) | f | $t_{ex}$ (ms) | $D_i$ (μm$^2$/ms) | $D_e$ (μm$^2$/ms) |
|---|---|---|---|---|---|---|
| PC-L | 7.0 ± 0.5 | 6.6 ± 0.4 | 0.50 ± 0.03 | 24.2 ± 4.1 | 3.09 ± 0.05 | 1.56 ± 0.19 |
| PC-R | 7.2 ± 0.5 | 6.8 ± 0.5 | 0.49 ± 0.03 | 28.0 ± 7.0 | 3.02 ± 0.08 | 1.59 ± 0.24 |
| PreCG-L | 7.1 ± 0.4 | 8.0 ± 0.4 | 0.47 ± 0.02 | 20.9 ± 2.2 | 2.89 ± 0.08 | 1.07 ± 0.10 |
| PreCG-R | 7.0 ± 0.4 | 8.0 ± 0.6 | 0.49 ± 0.01 | 21.6 ± 2.7 | 2.94 ± 0.06 | 1.11 ± 0.11 |
| PCG-L | 7.0 ± 0.5 | 7.8 ± 0.4 | 0.47 ± 0.02 | 21.4 ± 3.5 | 2.94 ± 0.06 | 1.22 ± 0.13 |
| PCG-R | 7.0 ± 0.5 | 7.8 ± 0.6 | 0.49 ± 0.02 | 20.9 ± 3.3 | 2.98 ± 0.07 | 1.23 ± 0.16 |
| CUN-L | 7.2 ± 0.5 | 7.1 ± 0.5 | 0.49 ± 0.01 | 19.7 ± 1.9 | 3.08 ± 0.09 | 1.36 ± 0.17 |
| CUN-R | 7.5 ± 0.5 | 7.4 ± 0.4 | 0.50 ± 0.02 | 23.0 ± 2.4 | 3.07 ± 0.07 | 1.33 ± 0.17 |
| STG-L | 6.9 ± 0.3 | 6.9 ± 0.1 | 0.40 ± 0.01 | 17.9 ± 0.7 | 2.85 ± 0.11 | 0.98 ± 0.05 |
| STG-R | 7.2 ± 0.4 | 6.9 ± 0.2 | 0.41 ± 0.01 | 17.3 ± 0.4 | 2.88 ± 0.09 | 0.99 ± 0.07 |
| MTG-L | 6.7 ± 0.3 | 7.1 ± 0.2 | 0.39 ± 0.02 | 17.1 ± 1.1 | 2.93 ± 0.08 | 0.92 ± 0.06 |
| MTG-R | 6.9 ± 0.5 | 7.1 ± 0.3 | 0.40 ± 0.01 | 17.4 ± 1.4 | 2.89 ± 0.08 | 0.90 ± 0.03 |
| IPL-L | 6.7 ± 0.6 | 7.5 ± 0.4 | 0.45 ± 0.02 | 17.9 ± 1.8 | 3.02 ± 0.11 | 1.02 ± 0.12 |
| IPL-R | 7.1 ± 0.5 | 7.6 ± 0.4 | 0.45 ± 0.02 | 17.5 ± 0.8 | 2.96 ± 0.13 | 0.99 ± 0.10 |
| SFG-L | 6.0 ± 0.4 | 7.4 ± 0.3 | 0.43 ± 0.02 | 18.2 ± 1.6 | 2.86 ± 0.11 | 1.00 ± 0.08 |
| SFG-R | 6.1 ± 0.5 | 7.6 ± 0.4 | 0.44 ± 0.02 | 19.0 ± 2.3 | 2.91 ± 0.08 | 0.95 ± 0.06 |
| **Cortical ribbon** | 7.0 ± 0.6 | 7.3 ± 0.5 | 0.43 ± 0.05 | 21.6 ± 4.9 | 2.83 ± 0.21 | 1.07 ± 0.15 |

**Table S3. Pearson correlation analysis results for pairs of ihMTR and SMI-derived metrics in white matter tracts:** f – axonal water fraction, $D_a$ – axonal water diffusivity, $D_e^{\parallel}$ – extracellular parallel diffusivity, $D_e^{\perp}$ –extracellular perpendicular diffusivity, $p_2$ – axonal dispersion. Pearson's correlation coefficient (r) is indicated along with the confidence interval (CI) and statistical significance (p-val) before and after False Discovery Rate (FDR) correction for multiple comparisons.

| **ihMTR vs.** | **f** | $D_a$ | $D_e^{\parallel}$ | $D_e^{\perp}$ | $p_2$ |
|---|---|---|---|---|---|
| **Pearson's r [CI]** | **0.74 [0.57 0.85]** | **0.41 [0.14 0.62]** | **0.66 [0.46 0.8]** | 0.25 [-0.04 0.5] | 0.03 [-0.26 0.31] |
| **p-val** | 2.89E-09 | 4.15E-03 | 4.43E-07 | 8.96E-02 | 8.47E-01 |
| **p-val FDR corrected** | 2.70E-08 | 8.93E-03 | 3.10E-06 | 1.32E-01 | 8.78E-01 |

**Table S4. Pearson correlation analysis results for pairs of MPF and SMI-derived metrics in white matter tracts:** f – axonal water fraction, $D_a$ – axonal water diffusivity, $D_e^{\parallel}$ –extracellular parallel diffusivity, $D_e^{\perp}$ – extracellular perpendicular diffusivity, $p_2$ – axonal dispersion. Pearson's correlation coefficient (r) is indicated along with the confidence interval (CI) and statistical significance (p-val) before and after False Discovery Rate (FDR) correction for multiple comparisons.

| **MPF vs.** | **f** | $D_a$ | $D_e^{\parallel}$ | $D_e^{\perp}$ | $p_2$ |
|---|---|---|---|---|---|
| **Pearson's r [CI]** | **0.57 [0.34 0.74]** | **0.44 [0.17 0.64]** | **0.65 [0.44 0.79]** | **0.48 [0.22 0.67]** | -0.13 [-0.4 0.16] |
| **p-val** | 2.91E-05 | 2.23E-03 | 8.66E-07 | 6.27E-04 | 3.84E-01 |
| **p-val FDR corrected** | 1.02E-04 | 5.67E-03 | 4.85E-06 | 1.95E-03 | 4.85E-01 |

**Table S5. Pearson correlation analysis results for pairs of ihMTR and SMEX-derived metrics in cortical gray matter ribbon:** f – cell-process fraction, $t_{ex}$ – exchange time, $D_e$ – extra-neurite diffusivity, $D_i$ – intra-neurite diffusivity. Pearson's correlation coefficient (r) is indicated along with the confidence interval (CI) and statistical significance (p-val) before and after False Discovery Rate (FDR) correction for multiple comparisons.

| ihMTR vs. | f | $t_{ex}$ | $D_e$ | $D_i$ |
|---|---|---|---|---|
| **Pearson's r [CI]** | -0.002 [-0.25 0.25] | **0.57 [0.37 0.72]** | 0.27 [0.02 0.49] | -0.12 [-0.36 0.13] |
| **p-val** | 9.85E-01 | 1.25E-06 | 3.54E-02 | 3.54E-01 |
| **p-val FDR corrected** | 9.85E-01 | 6.23E-06 | 5.32E-02 | 4.41E-01 |

**Table S6. Pearson correlation analysis results for pairs of MPF and SMEX-derived metrics in cortical gray matter ribbon:** f – cell-process fraction, $t_{ex}$ – exchange time, $D_e$ – extra-neurite diffusivity, $D_i$ – intra-neurite diffusivity. Pearson's correlation coefficient (r) is indicated along with the confidence interval (CI) and statistical significance (p-val) before and after False Discovery Rate (FDR) correction for multiple comparisons.

| MPF vs. | f | $t_{ex}$ | $D_e$ | $D_i$ |
|---|---|---|---|---|
| **Pearson's r [CI]** | **0.49 [0.27 0.66]** | 0.07 [-0.18 0.32] | -0.11 [-0.35 0.14] | 0.27 [0.03 0.49] |
| **p-val** | 5.74E-05 | 5.69E-01 | 3.83E-01 | 3.06E-02 |
| **p-val FDR corrected** | 1.72E-04 | 6.10E-01 | 4.41E-01 | 5.11E-02 |

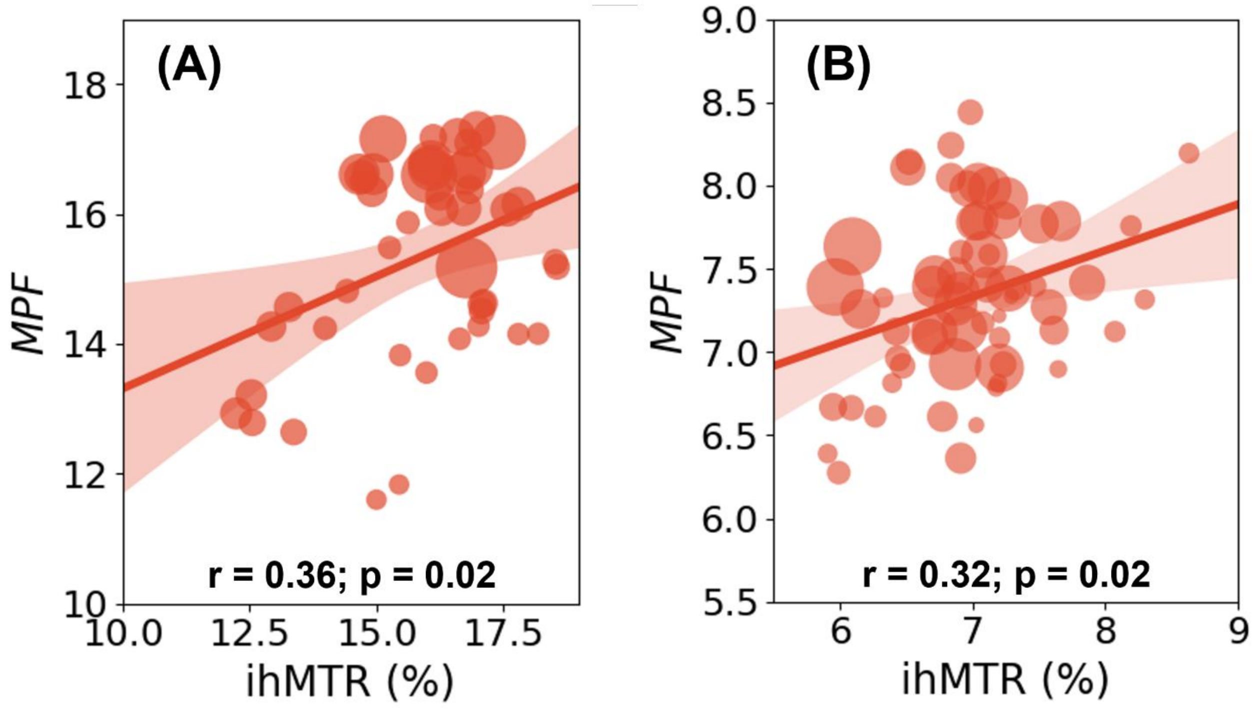


**Figure S1.** Correlations between ihMTR and MPF in (A) WM and (B) GM brain regions. Linear regression lines with 95% confidence intervals are displayed for visual guidance. Pearson's correlation coefficients (r) and corresponding p-values (p) FDR-corrected for multiple comparisons are indicated in each plot; values shown in **bold** denote significant correlations after FDR correction.

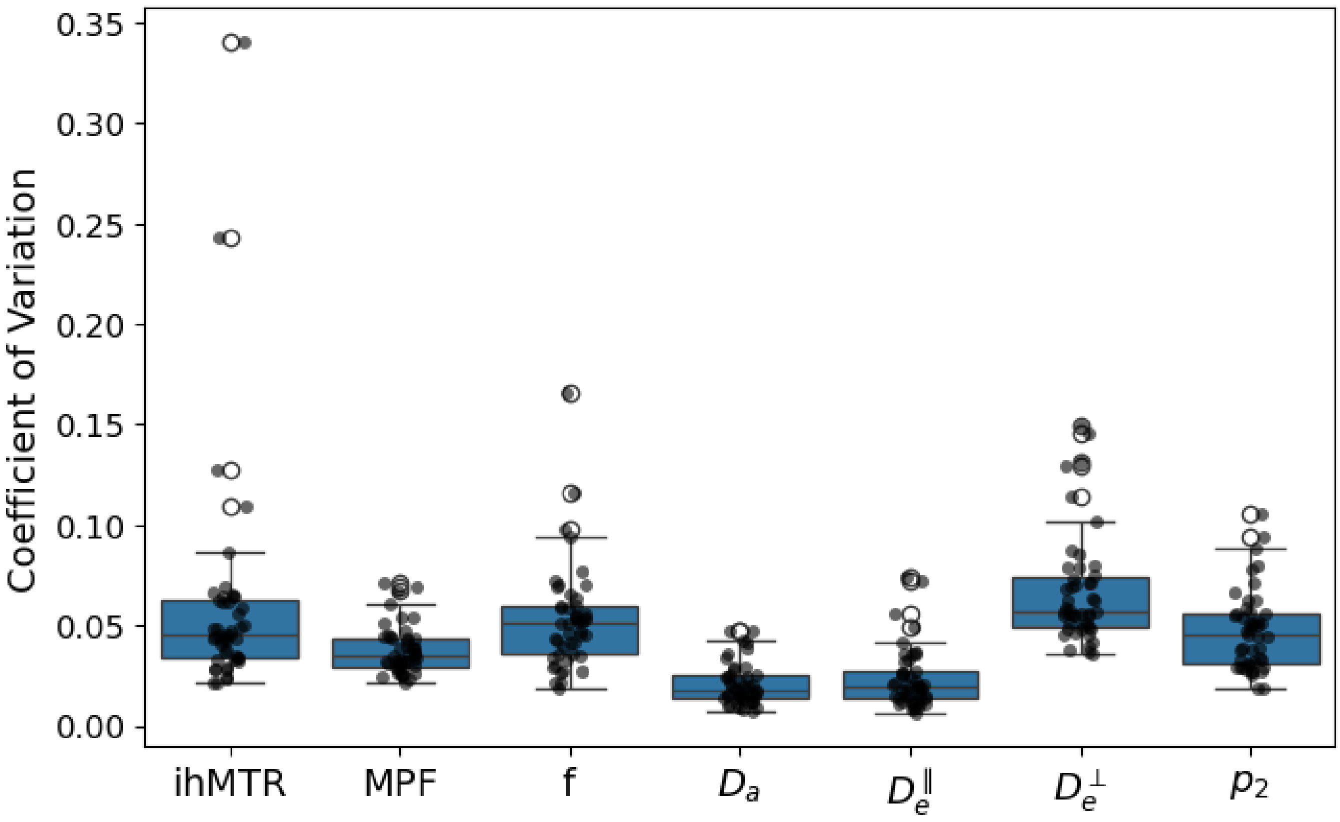


**Figure S2.** Coefficient of variation for ihMTR, MPF and SMI metrics, calculated as the standard deviation of the mean value across subject divided by the mean of the mean values across subjects for each white matter tract.

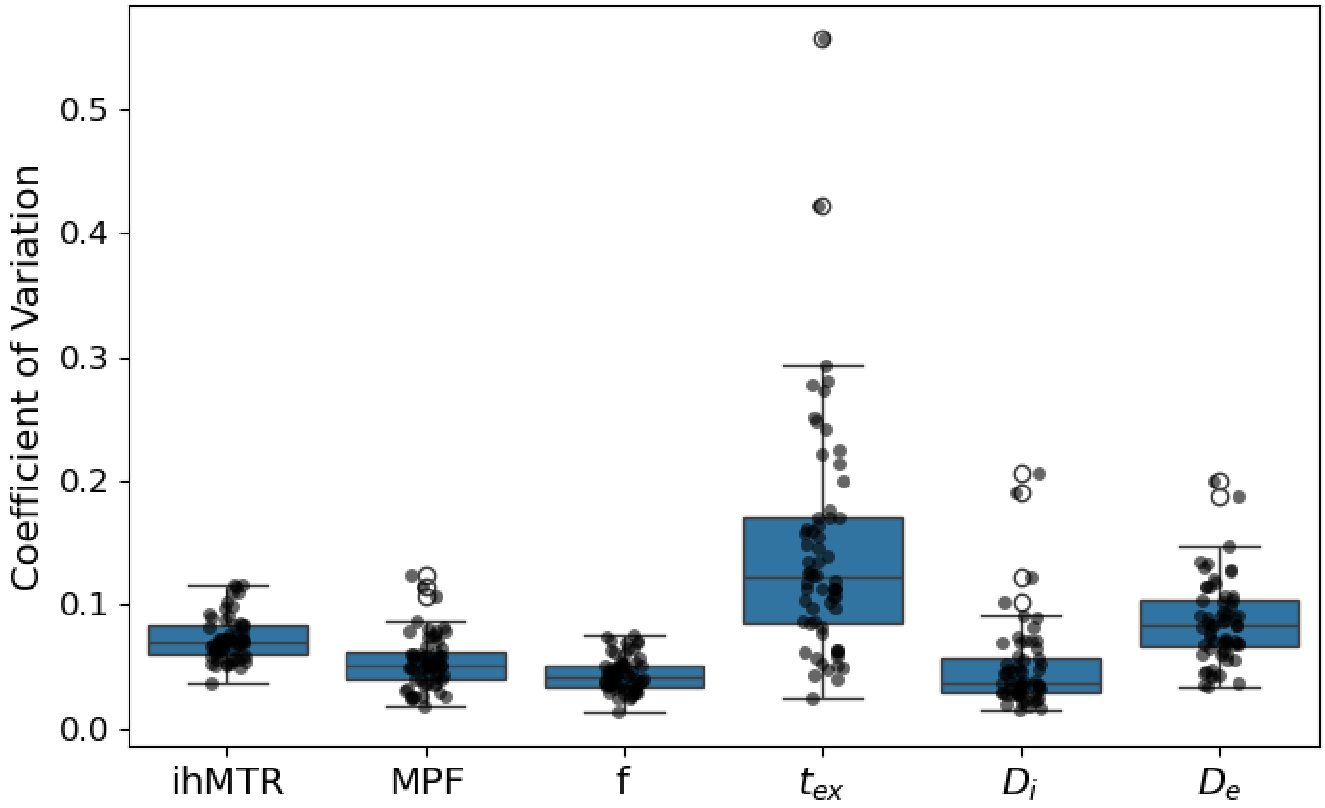


**Figure S3.** Coefficient of variation for ihMTR, MPF and SMEX metrics, calculated as the standard deviation of the mean value across subject divided by the mean of the mean values across subjects for gray matter ribbon region.

.
